# Neanderthal Introgressed COVID-19 Risk Locus 3p21.31 Under Positive Selection in East Asia via LZTFL1-Driven Ciliary Changes

**DOI:** 10.1101/2025.10.01.679818

**Authors:** Rudra Kumar Pandey, Shailesh Desai, Gyaneshwer Chaubey

**Affiliations:** Gyan Lab, Cytogenetics Laboratory, Department of Zoology, Banaras Hindu University, Varanasi, India-221005

**Author notes:** Corresponding authors (Rudra Kumar Pandey), (Gyaneshwer Chaubey).

**Keywords:** COVID-19, 3p21.31, East Asia, Positive Selection, LZTFL1

## Abstract

The Neanderthal introgressed 3p21.31 locus has been repeatedly associated as a major host genetic determinant of severe COVID-19, particularly in European and South Asian populations. However, the evolutionary history and likely effector gene remain unclear. We investigated signatures of historical selection at this locus and the mechanism by which it might influence COVID-19 severity. Using 1000 Genomes Project data from diverse global populations, we applied various selection scans methods and integrated it with functional analysis. Multiple, independent lines of evidence indicate strong positive selection at 3p21.31 in East Asians. The region exceeds the 99.99th-percentile PBS threshold in EAS, shows extended ROH and the slowest LD decay among populations, and exhibits reduced nucleotide diversity and significantly negative Tajima’s D, consistent with a selective sweep. XP-EHH identified a pronounced signal centred on LZTFL1. Marginal-tree reconstruction indicates the derived allele arose in modern humans and swept to high frequency in East Asia at a time consistent with an ancient corona or related virus-like epidemic ∼20,000-25,000 (20-25 Kya) years ago. LD and pathway analyses implicate a high-LD haplotype tagging LZTFL1 shows significant enrichment for cilium-assembly and BBSome-mediated cargo-targeting pathways likely causing subtle shifts in cilium number or beat frequency, in lung and airway epithelia, thereby profoundly affecting the mucus and pathogens clearance. Based on the Overall observation, we propose that after the Neanderthal introgression, East Asia went through pathogen pressure for ∼20-25 Kya old ancient coronavirus- or related virus epidemic leading to positive selection at 3p21.31locus. These findings underscore the complex interplay between ancient admixture and adaptive events linked to modern COVID-19 severity.

## Introduction

The Coronavirus disease 2019 (COVID-19), caused by a positive novel single-stranded RNA Virus known as severe acute respiratory syndrome coronavirus 2 (SARS-CoV-2) has caused the deaths of over 18 million people to date and has posed significant challenges to both the healthcare system and socioeconomic status worldwide (Zhou et al. 2020).

There is significant diversity in the symptoms experienced by patients infected with SARS-CoV-2. These symptoms can range from asymptomatic to mild and severe illness, with severity often increasing with age and comorbidities (Russell, Lone, and Baillie 2023). However, not all individuals with severe COVID-19 symptoms have these known risk factors, as many young and healthy people have also been severely impacted. In this context, numerous studies have been conducted across different populations to identify the link between genomic variants and the severity of COVID-19 (Pandey et al. 2021, 2024; Srivastava, Bandopadhyay, et al. 2020; Srivastava, Pandey, et al. 2020).

A landmark Genome Wide Association study (GWAS) conducted during the early stages of the pandemic identified a cluster of genes (SLC6A20, LZTFL1, CCR9, FYCO1, CXCR6, and XCR1) at the 3p21.31 locus that were strongly linked to severe COVID-19 outcomes, as it can nearly double the likelihood of experiencing severe COVID-19 (null 2020). This association remained the most significant and replicated, particularly in individuals of European (EUR) and South Asian (SAS) ancestries (Kanai et al. 2023; Niemi et al. 2021; Pairo-Castineira et al. 2023; Wu et al. 2021). Despite the presence of numerous genes at the 3p21.31 locus, pinpointing a single causal gene directly contributing to the pathogenesis of COVID-19 proved to be difficult (null 2020).

Beyond its role in influencing the severity of COVID-19, the 3p21.31 locus is also connected to several other health conditions, including type 1 diabetes, developmental delays, elevated serum creatine kinase levels, and heart conditions (Eto et al. 2013; Nikpay and McPherson 2021; Tran et al. 2021).

The evolutionary history of the 3p21.31 locus provides an additional intrigue. It has been proposed that this region harbours a haplotype inherited from Neanderthals. The presence of this archaic haplotype in contemporary populations varies significantly across geographic regions, suggesting that historical selective pressure may have shaped its distribution (Zeberg and Pääbo 2020). These findings underscore the complex interplay between ancient admixture events and susceptibility to modern human diseases. However, the nature of these pressures, whether driven by infectious diseases, environmental factors, or other evolutionary forces, remains an open question.

Therefore, understanding the selection dynamics acting on the 3p21.31 locus is crucial for unravelling the mechanisms underlying its association with COVID-19. Investigating whether these genes have been targets of selection in human populations can shed light on their evolutionary significance and potential roles in conferring disease resistance or susceptibility.

Reflecting on the current evidence concerning the association of the 3p21.31 locus with COVID-19 severity, alongside the influence of the Neanderthal genome, this study aimed to examine the selective history of the 3p21.31 locus, leveraging a combination of population genetics tools and comparative genomic approaches. By exploring signatures of selection and functional consequences of genetic variation in this region, we sought to elucidate the evolutionary forces as well as the mechanism by which it might influence COVID-19 severity or susceptibility, which may provide valuable insights into its role in ancient selection events in contemporary pandemics and inform future therapeutic strategies for infectious diseases.

## Result and Discussion

### Positive selection at 3p21.31 in East Asians

Our detailed Population Branch Statistic (PBS) analysis highlighted the 3p21.31 locus as a strong candidate for positive selection in East Asians (EAS). This region exhibited a highly significant PBS value, exceeding the significance threshold (99.99% percentile), which indicates pronounced genetic differentiation at the 3p21.31 locus between the EAS and other populations (Figure 1). The elevation of the PBS value at 3p21.31 signifies that this region might have undergone unique evolutionary pressures in the EAS, suggesting a localised selection event. Such selection is likely driven by environmental or pathogenic factors specific to EAS, where variants in this region may have provided a selective advantage.

**Figure 1.**
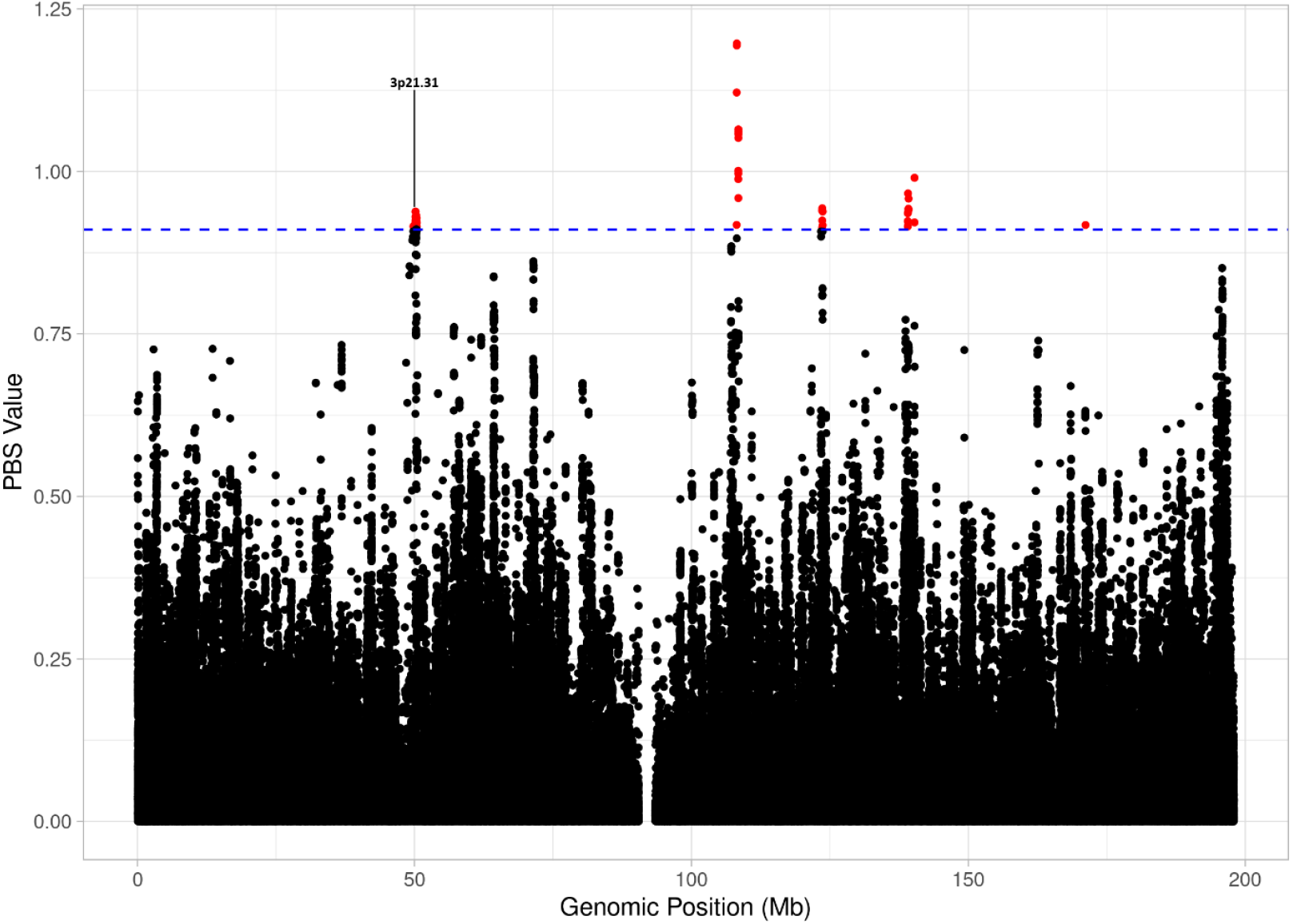
PBS scan of EAS: The plot represents PBS values across the 3^rd^ chromosome (genomic position in Mb) comparing EAS to other populations. Each point is a tested variant; the horizontal dashed blue line denotes the 99.99th-percentile empirical significance threshold. Red points exceed this threshold and represent highly differentiated loci. A prominent peak at the 3p21.31 region (annotated) rises well above the significance cutoff, indicating pronounced genetic differentiation consistent with a localized selection signal in EAS.

### 3p21.31 shows extended homozygosity in East Asians

The Runs of Homozygosity (ROH) analysis provides additional evidence of positive selection at the 3p21.31 locus. EAS displays the longest mean ROH segment length and the highest mean number of segments among the studied populations (Figure 2). The extended ROH length and increased segment count indicate significant homozygosity over large genomic regions in EAS, compared to other populations, a hallmark of selective sweeps. Such patterns arise when advantageous alleles rapidly increase in frequency, reducing genetic variation in the surrounding genomic regions. This supports the notion that strong selection has shaped this locus in EAS, potentially in response to regional environmental challenges or disease exposure.

**Fig 2.**
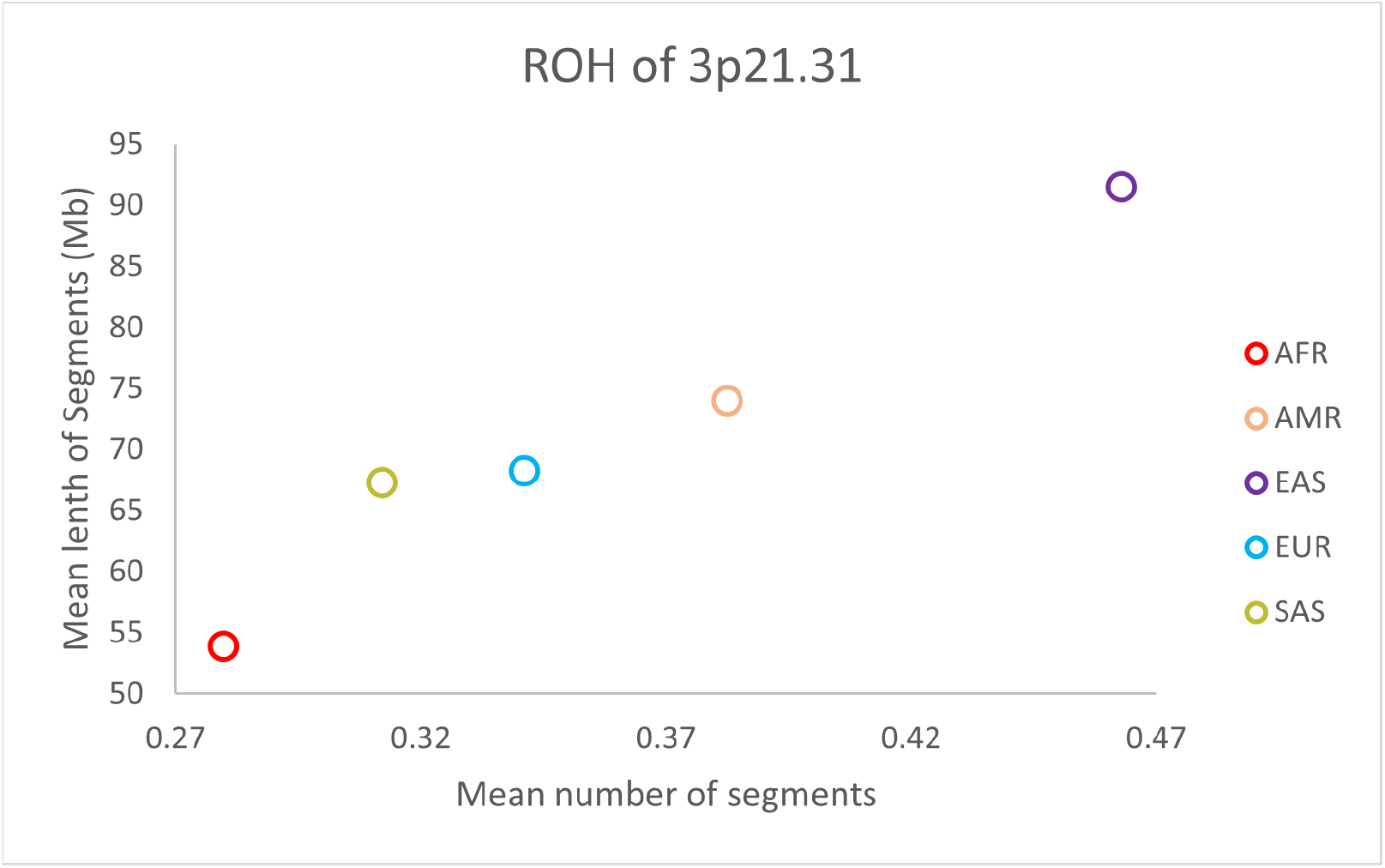
ROH at 3p21.31 across continental populations: The Scatterplot represents mean ROH segment count (x-axis) versus mean ROH segment length in megabases (y-axis) calculated for the 3p21.31 locus in five continental groups: AFR, AMR, EAS, EUR and SAS. The EAS show both the highest mean number of ROH segments and the longest mean segment length, indicating markedly extended homozygosity at 3p21.31 relative to other populations.

### Lower molecular diversity at 3p21.31 in EAS

The molecular diversity indices also highlighted reduced genetic diversity at the 3p 21.31 locus in the EAS populations. EAS exhibited the lowest nucleotide diversity (π) among all studied populations, indicating a depletion of overall genetic variation at this locus. Additionally, other diversity metrics, such as θ_k_ (coalescent-based diversity), were also diminished in EAS compared with other populations (Figure 3). This reduced diversity reflects the genetic homogenisation associated with positive selection, where advantageous alleles sweep through the population, reducing variability in the surrounding regions.

**Figure 3.**
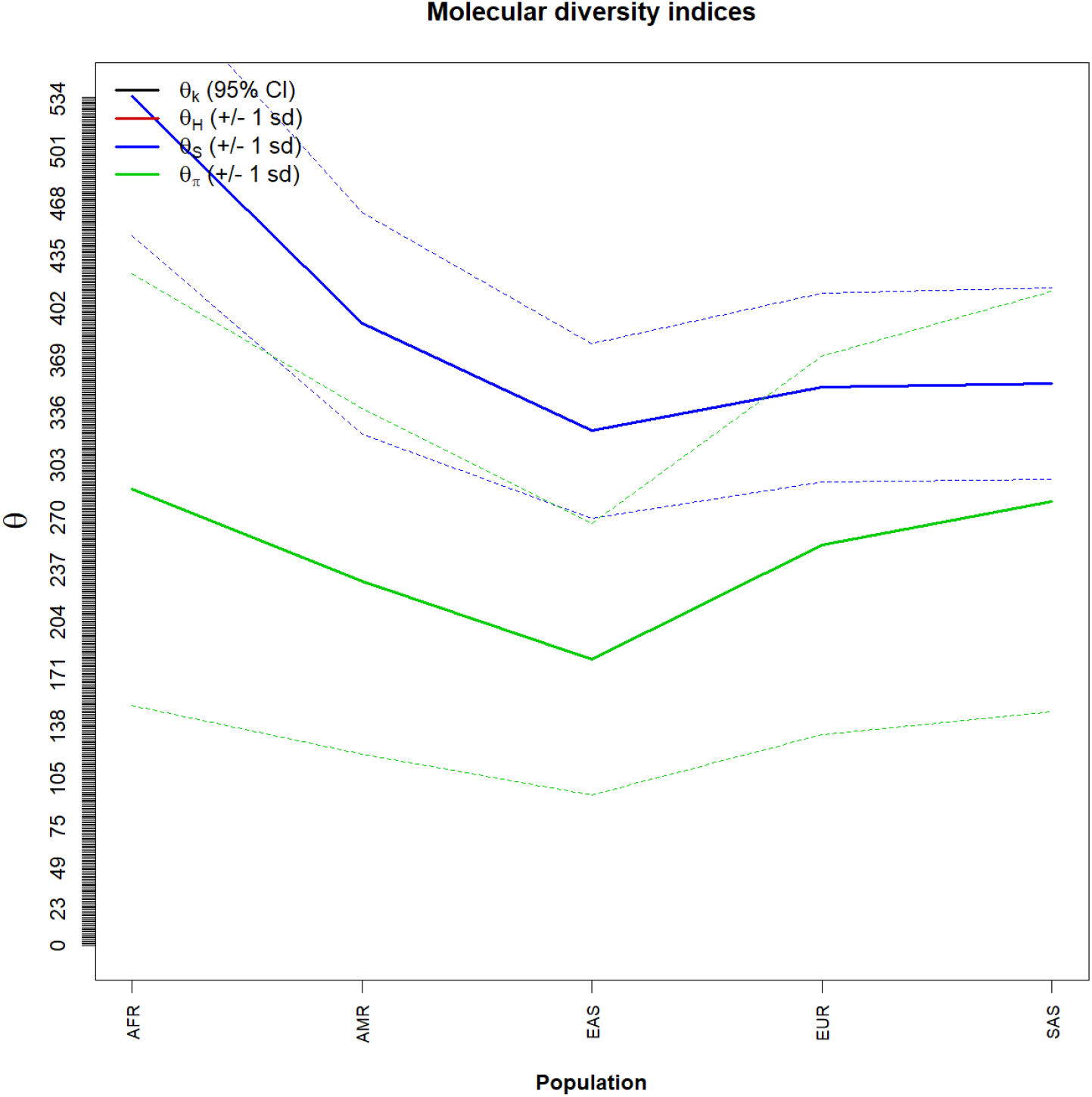
Molecular diversity indices at 3p21.31 across continental populations: Line plot of molecular diversity indices calculated for the 3p21.31 locus in five continental groups (AFR, AMR, EAS, EUR, SAS). The plotted statistics are: θ_k_ (coalescent-based diversity; 95% CI), and θπ (nucleotide diversity; ±1 SD). Each point on the x-axis corresponds to the population mean with their uncertainty bands (95% CI or ±1 SD). East Asians (EAS) display the lowest nucleotide diversity (π) and θ_k_ relative to other populations, indicating a depletion of standing variation at 3p21.31, consistent with reduced genetic variation at this locus.

### Low pairwise differences at 3p21.31 in EAS

The heatmap of the average number of pairwise differences provided further context for the patterns observed at the 3p21.31 locus. Within the EAS population, the pairwise differences were lowest than those observed in other populations, indicating reduced intra-population diversity. Similarly, the between-population differences involving EAS are distinct, reflecting genetic differentiation driven by selection at this locus. The low pairwise differences within the EAS emphasise the reduced variability resulting from the fixation of advantageous alleles, whereas the distinct differences between the EAS and other populations highlight the population-specific evolutionary pressures acting on the 3p21.31 locus (Figure 4).

**Figure 4.**
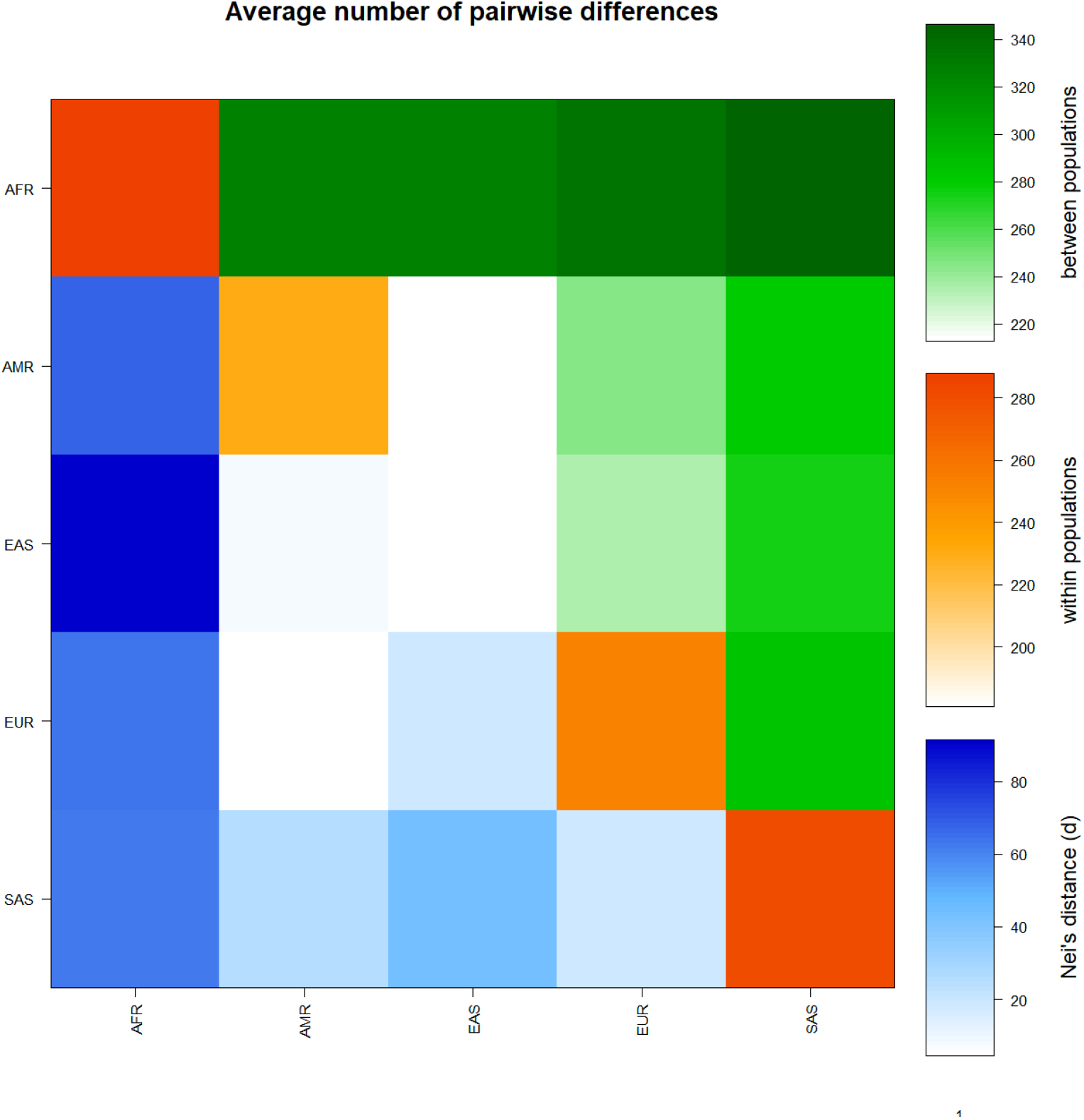
Heatmap of the average number of pairwise differences at 3p21.31 across populations: Heat map showing intra- and inter-population variation measured by average pairwise sequence differences at the 3p21.31 locus. The average pairwise differences between populations are shown in the upper triangle of the matrix (green). The average number of pairwise differences within each population group are shown along the diagonal (orange). The differences between populations based on Nei’s genetic distances are depicted in the lower triangle of the matrix (blue). The obtained values of various parameters are shown on the colour scales. The EAS population exhibits markedly lower within-population pairwise differences than the other groups, indicating reduced intra-population diversity at this locus, consistent with partial or complete fixation of an advantageous allele. Between-population comparisons that include EAS are similarly distinct from other pairings, reflecting increased genetic differentiation of EAS at 3p21.31.

### Signatures of non-neutral evolution at 3p21.31 in EAS

Tajima’s D analysis provided critical insights into the demographic and selective history of the 3p21.31 locus in EAS populations. EAS exhibited a significantly negative Tajima’s D value for this region, indicating an excess of low-frequency polymorphisms, suggesting that recent positive selection or population expansion has reduced the proportion of intermediate-frequency alleles in this region. Moreover, the significant p-value associated with Tajima’s D further supports the hypothesis of non-neutral evolutionary processes (Table 1).

**Table 1.**
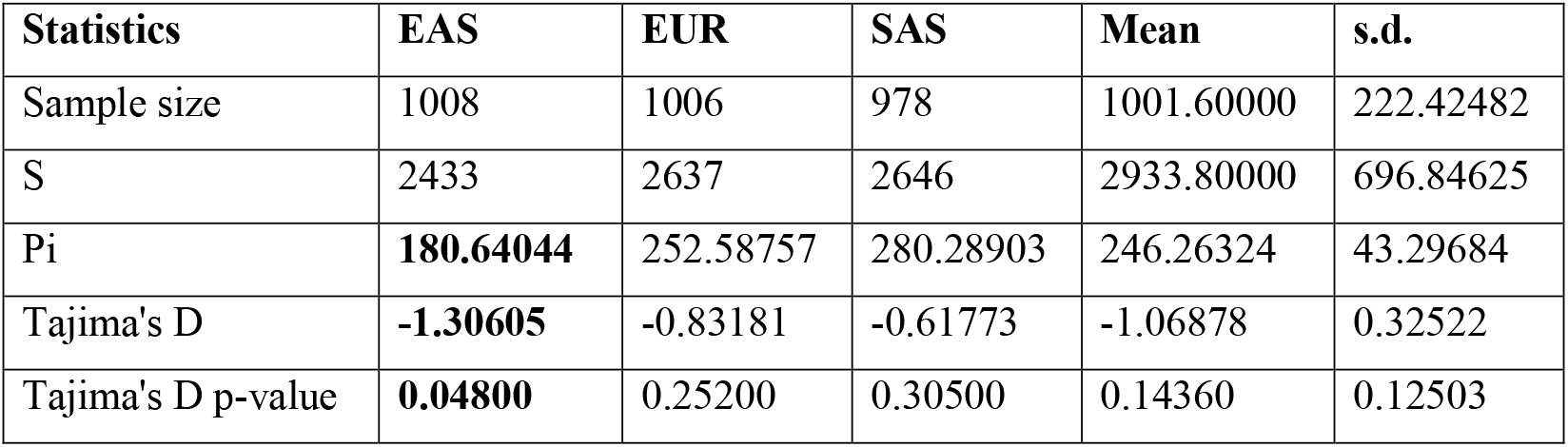
Tajima’s D results and diversity statistics at 3p21.31: The Tajima’s D neutrality test results for populations (EAS, EUR, SAS), showing sample size, number of segregating sites (S), nucleotide diversity (π), Tajima’s D values, and corresponding p-values. These results indicate deviations from neutral evolution, with significant Tajima’s D p-values (e.g., EAS, 0.048) indicating potential demographic or selective events.

### Slower LD decay at 3p21.31 in EAS

The analysis of Linkage Disequilibrium (LD) decay at the 3p21.31 locus across five 1000 Genome populations such as African (AFR), American (AMR), EAS, EUR, and SAS indicates distinct patterns of LD decline over increasing genomic distances (Figure 5). The EAS population exhibited the highest initial r^2^ values and the slowest LD decay, indicating a stronger and more extended linkage disequilibrium, which suggests reduced recombination rates associated with recent positive selection at this locus. In contrast, the African population demonstrated the lowest r^2^ values and the most rapid LD decay, a pattern typically associated with higher genetic diversity and recombination rates. The EUR and SAS populations showed an intermediate LD decay pattern, with r^2^ values decreasing more rapidly than in EAS but remaining higher than in Africans over longer distances. These differences in LD decay dynamics suggest population-specific selection pressures and recombination landscapes at 3p21.31, with a potential signal of stronger selection in East Asians.

**Figure 5.**
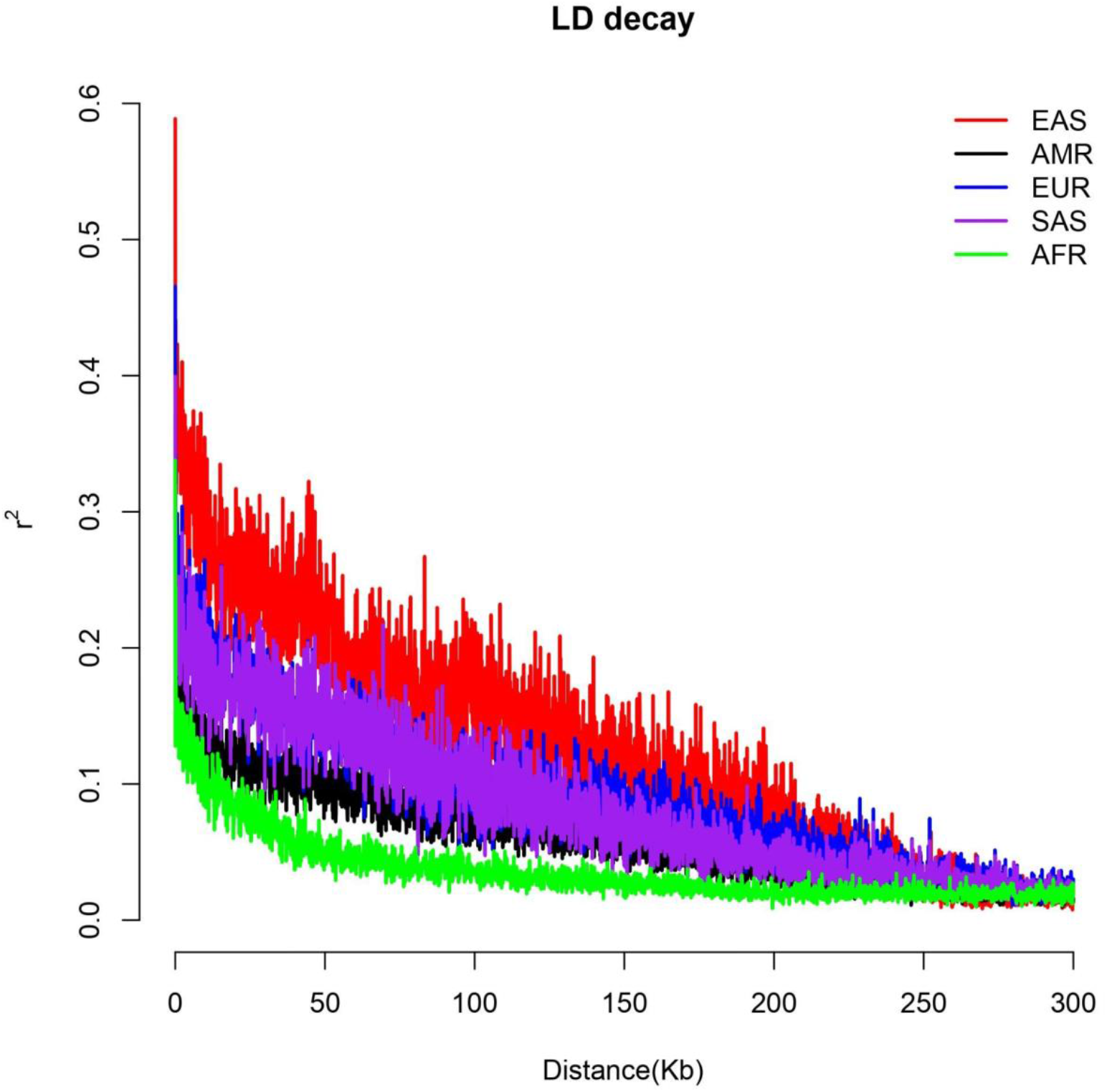
Comparative LD decay profiles of the 3p21.31 locus across global populations: The plot shows the decline in pairwise LD (measured as r^2^, y-axis) with increasing physical distance between SNPs (in kilobases, x-axis) in continental populations. The colours correspond to populations as shown in the legend (EAS = red, AMR = black, EUR = blue, SAS = purple, AFR = green). The EAS curve exhibits the highest initial r^2^ and the slowest decay with distance, indicating more extended linkage disequilibrium consistent with reduced local recombination or recent positive selection at this locus.

### Selection signal at LZTFL1 in EAS

The Cross-population extended haplotype homozygosity (XP-EHH) analysis was conducted using EUR as the reference population and EAS as the target population at the 3p21.31 locus. The results showed significantly high positive XP-EHH scores exceeding the +2-standard deviation (SD) threshold, suggesting recent positive selection in the East Asian population, at SNP rs2064061, which overlaps with the LZTFL1 gene (Figure 6). The LZTFL1 gene has been previously identified as an effector gene at the COVID-19 risk locus, 3p21.31, associated with COVID-19 severity (Downes et al. 2021; Fink-Baldauf et al. 2022). The presence of extended haplotype homozygosity in the LZTFL1 gene suggests that beneficial alleles may have risen to high frequencies in EAS, possibly conferring an advantage in the immune response or other physiological traits.

**Figure 6.**
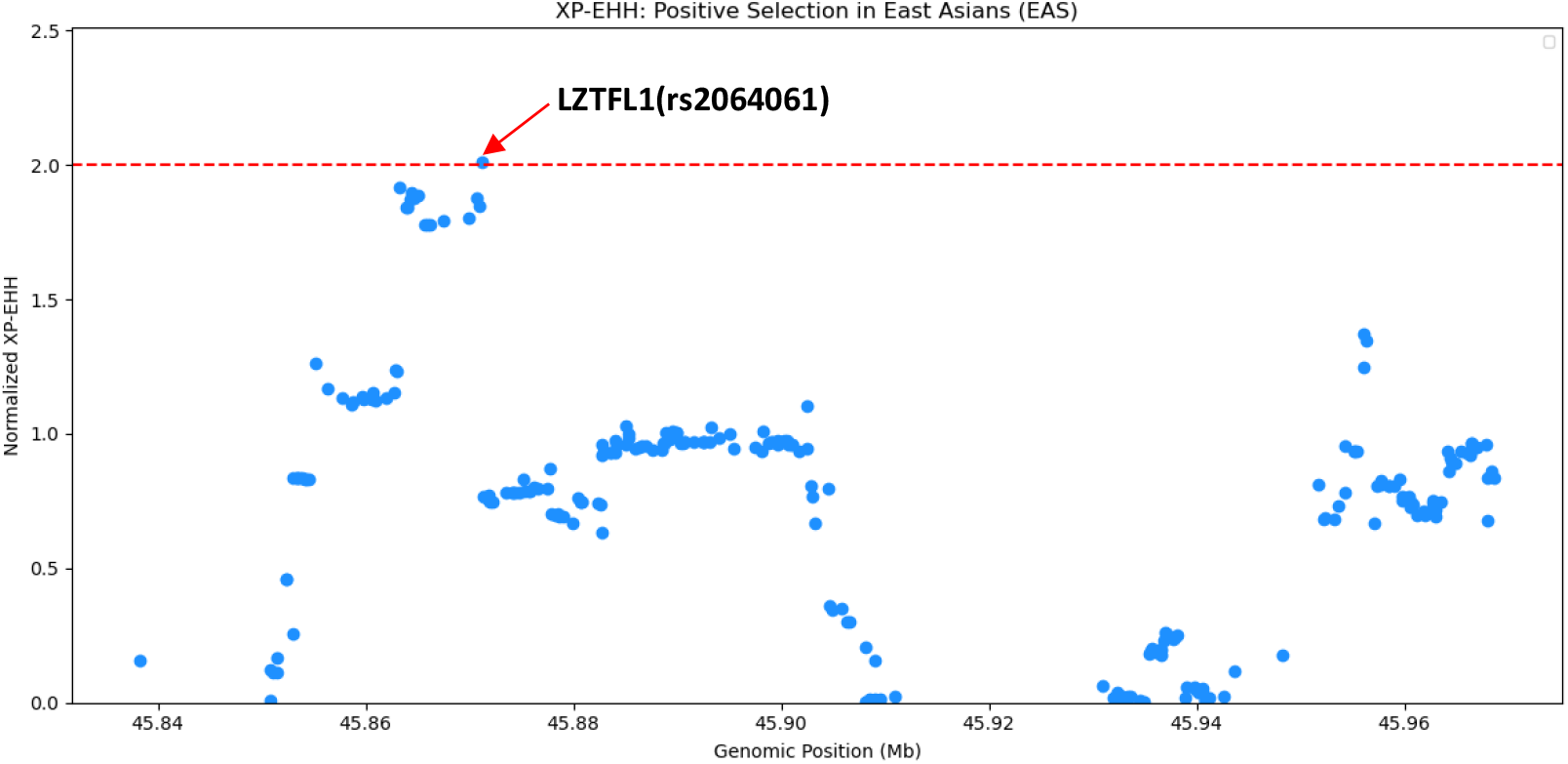
XP-EHH scan across the 3p21.31 locus in EAS: Each point represents an SNP’s normalised XP-EHH score plotted by genomic position (Mb); the red dashed line denotes the significance threshold (normalised XP-EHH = 2). A lead variant rs1058961 (LZTFL1) exceeds the threshold, indicating a strong signal of recent positive selection in the EAS population.

### Ancient Viral Epidemics as a Driver of LZTFL1 Selection in EAS

The Marginal-tree reconstruction for rs1058961 (LZTFL1) revealed a stark contrast between archaic and modern human lineages (Figure 7). All four archaic genomes (Altai, Chagyrskaya, Vindija Neanderthals, and Denisova) uniformly carry the ancestral allele, with no derived-allele branches observed. This absence confirms that the derived variant is exclusive to modern humans and must have originated after our lineage split from Neanderthals and Denisovans. In modern populations, we observed both ancestral and derived lineages, but at markedly different frequencies and branch length distributions. CHB (Han Chinese) samples showed the highest proportion of derived-allele branches. Independent analyses of EAS genomes suggest a corona or a closely related virus was epidemic in EAS ∼20,000-25,000 years ago (Souilmi et al. 2021). The timing of the elevated proportion of derived branches in CHB, is consistent with the timing of the epidemic, suggesting a strong pathogen-mediated selective sweep that elevated the derived allele to high frequency in EAS populations.

**Figure 7.**
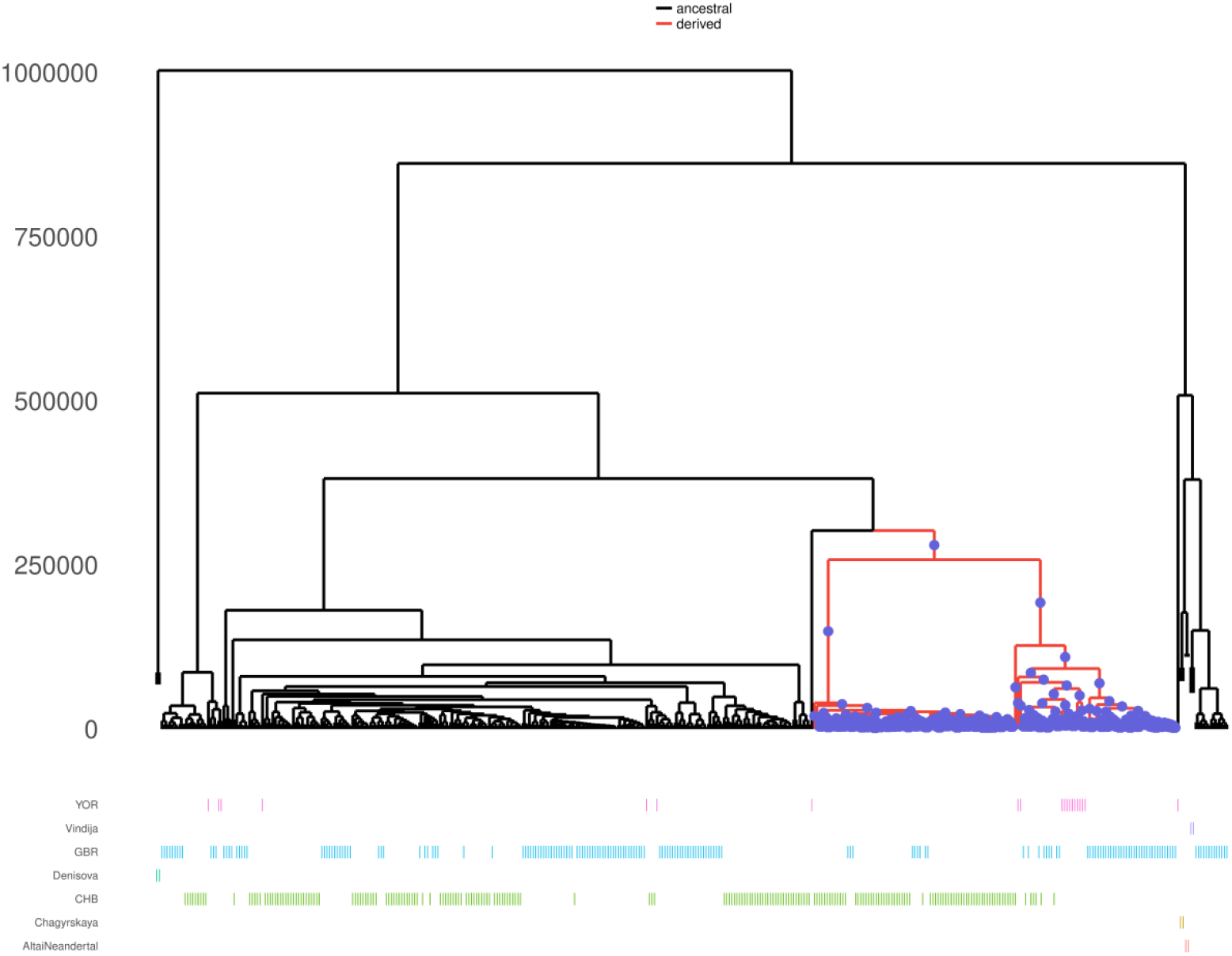
Marginal-tree reconstruction of rs1058961 (LZTFL1) in archaic and modern human populations: Branches carrying the ancestral allele are shown in black, and those carrying the derived allele in red. Tree depths (vertical axis) are scaled to the coalescence time. The colored bars below the tree map individuals from various archaic and modern human populations to their corresponding ancestral or derived alleles (YOR = Yoruba; GBR = Great Britain; CHB = Han Chinese; and the four archaic genomes: Altai, Chagyrskaya, Vindija Neanderthals; Denisova). All archaic genomes exclusively carry the ancestral allele (black) branches, indicating the derived variant is exclusive to modern humans and must have arisen after the split from Neanderthals/Denisovans. Modern samples show both ancestral and derived lineages with distinct branch-length and frequency patterns; CHB displays the highest proportion of derived-allele branches. The elevated proportion of derived branches in CHB is temporally consistent with an ∼20–25 kya viral epidemic event reported for EAS (Souilmi et al. 2021) and is compatible with a strong, pathogen-mediated selective sweep elevating the derived allele in EAS populations.

### Functional Adaptation via LZTFL1-Driven Ciliary Modulation

The LD analysis revealed that rs2064061 is part of a high-LD haplotype block (r^2^ > 0.8) encompassing five SNPs in the EAS population (rs113776340, rs9814578, rs9871972, rs11130076, and rs3172377). Pathway enrichment of the COVID-19–associated locus at LZTFL1 (marked by rs9871972) identified a significant overrepresentation of Cilium Assembly pathways, that is, BBSome-mediated cargo-targeting to cilium (pval=0.002135), cargo trafficking to the periciliary membrane (pval=0.004733), cilium assembly (pval=0.018747), and organelle biogenesis and maintenance (pval=0.027285) (Figure 8). This suggests that the protective allele of this SNP may contribute to resistance against progression to severe COVID-19 in EAS populations by causing subtle shifts in cilium number or beat frequency in lung and airway epithelia, thereby profoundly affecting mucus and pathogen clearance and limiting viral spread and inflammation (Downes et al. 2021; Fink-Baldauf et al. 2022; Seo et al. 2011; Song et al. 2024).

**Figure 8:**
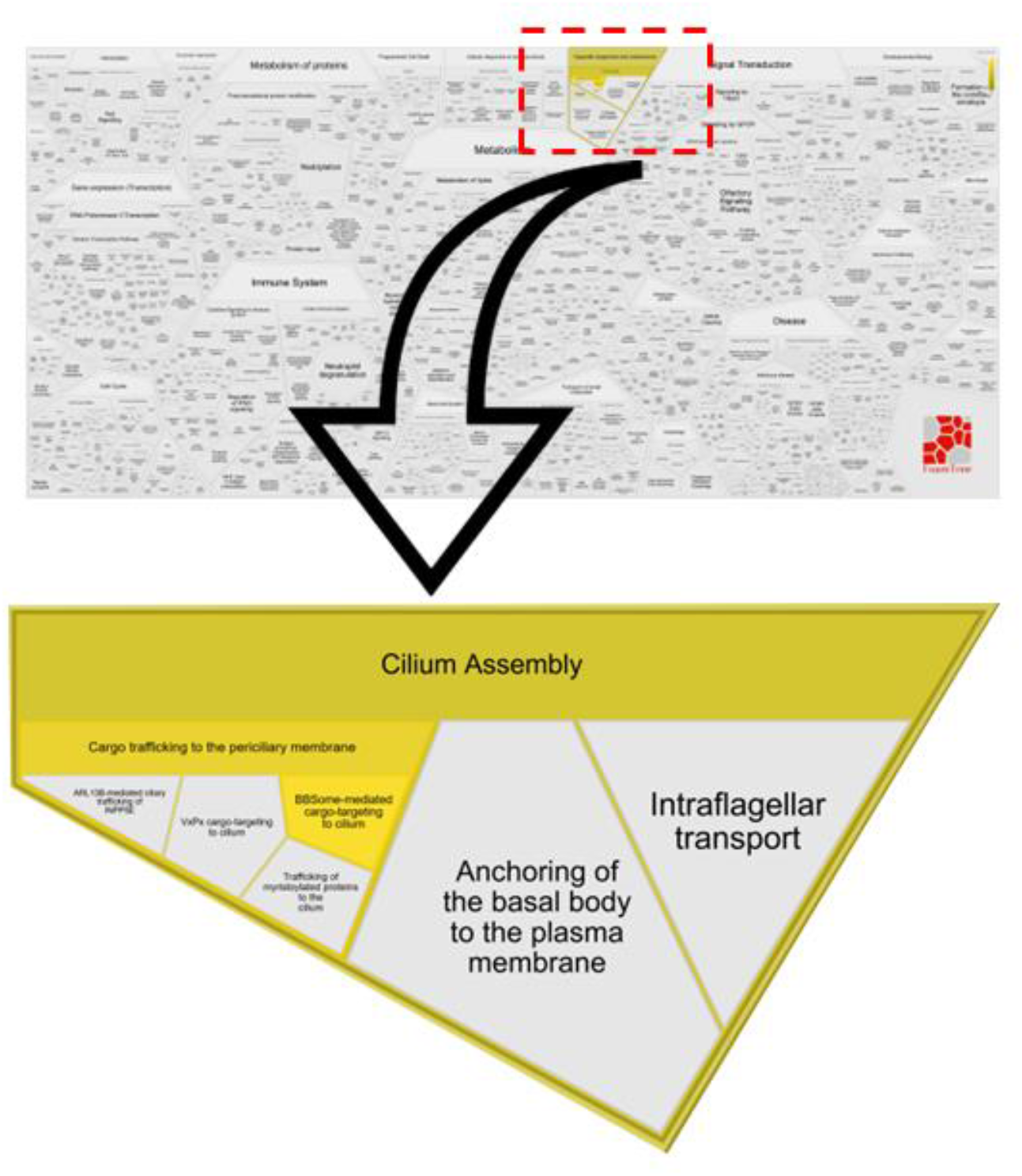
Pathway enrichment analysis of the COVID-19-associated locus at LZTFL1: The reactome pathway associated with rs9871972 (tagged by rs2064061) lies in a high-LD haplotype block (r^2^ > 0.8). The colour bar shows how the colour intensity represents the p-value of the statistical test for overrepresentation. The arrow indicates the location of the most significant zoomed sector highlighted by “Cilium Assembly pathways” suggesting that the protective allele carried on this EAS haplotype may alter ciliary number or beat properties in airway epithelia, with plausible consequences for mucus and pathogen clearance that could reduce progression to severe COVID-19.

### Evolutionary Shaping of LZTFL1 Associations Across Populations

Using the GENOMIC COVID-19 GWAS meta-analysis data (Pairo-Castineira et al. 2023), we observed that the 3p21.31 locus is significantly associated with severe COVID-19 in both European and South Asian cohorts. The SNP rs9871972 (LZTFL1) showed a highly significant protective effect (EUR: β = –0.19, p = 1.2 × 10^−19^; SAS: β = –0.37, p = 2.5 × 10^−9^), a pattern consistent with rs9871972 acting as a passenger SNP in strong linkage disequilibrium with the causal variant at 3p21.31 and carrying the protective allele on a broader haplotype. By contrast, the East Asian sample showed no significant association (β = –0.28, p = 0.063); nevertheless, the negative effect size in EAS is directionally consistent with a protective role (Figure 9). This lack of significance may reflect historical, population-specific selection at 3p21.31 in EAS potentially driven by an ancient coronavirus-like epidemic that altered haplotype structure and allele frequencies.

**Figure 9.**
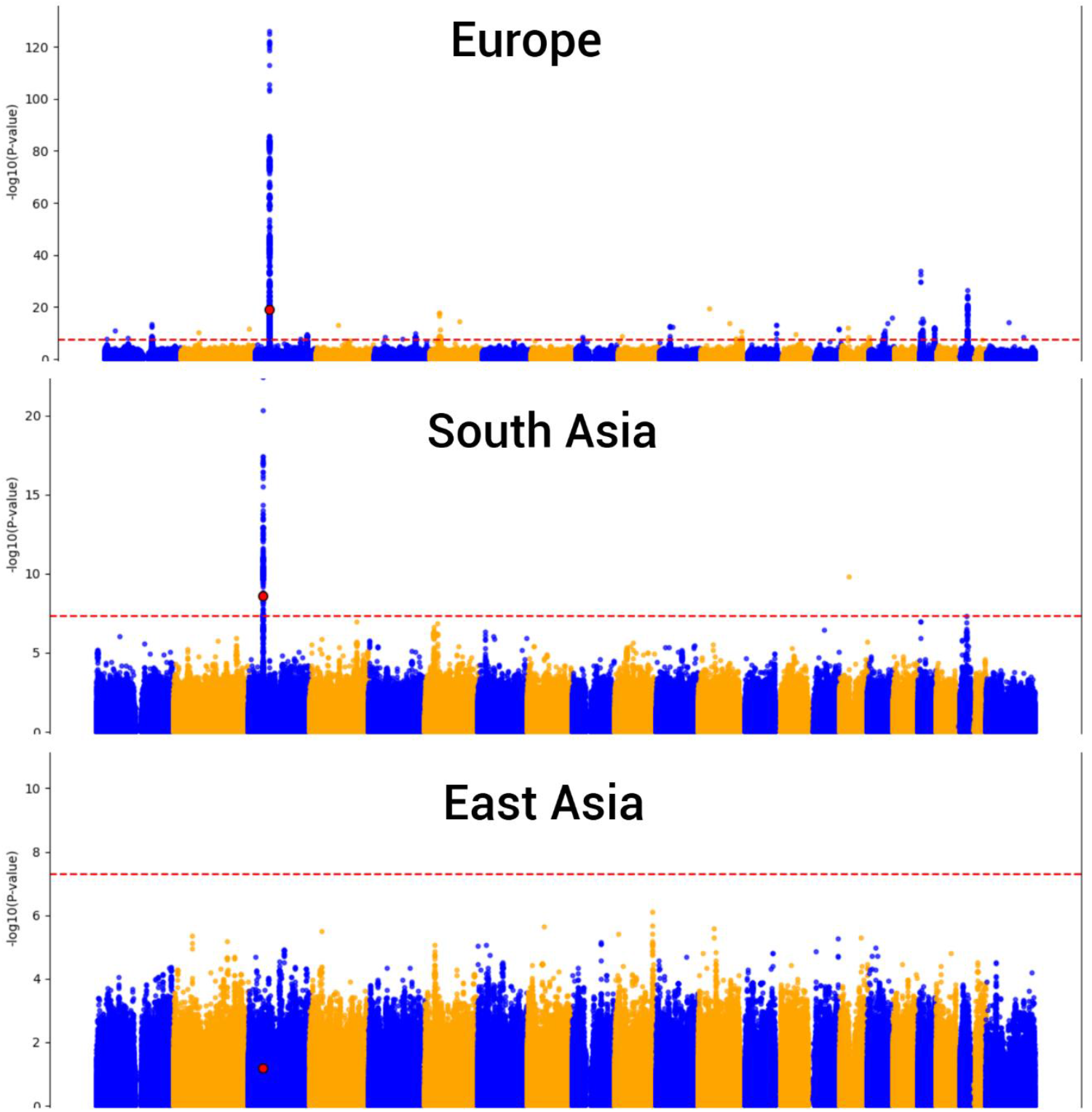
Ancestry-stratified COVID-19 GWAS meta-analysis highlighting the 3p21.31/LZTFL1 region: Top to bottom: Europe, South Asia, and East Asia. Points on the Y-axis are −log10(p-value) for tested variants (alternating chromosome colours for clarity); the horizontal dashed line marks the genome-wide significance threshold (p = 5×10^−8^).In the GENOMIC COVID-19 meta-analysis, the LZTFL1 variant rs9871972 at 3p21.31locus (highlighted in red) exhibits a strong protective effect in EUR (β = −0.19, p = 1.2×10^−19^) and SAS (β = −0.37, p = 2.5×10^−9^), whereas the EAS sample shows a non-significant but directionally consistent protective effect (β = −0.28, p = 0.063). These patterns are compatible with rs9871972 acting as a passenger SNP on a protective haplotype in EUR and SAS and with population-specific differences in haplotype architecture (e.g., a protective allele carried on a broad haplotype in EUR/SAS) and by historical selection at 3p21.31 in East Asia that may have altered allele frequencies and LD patterns in EAS.

## Conclusion

The 3p21.31 locus shows strong evidence of positive selection in East Asians, supported by multiple complementary lines of evidence. The high PBS values at the 3p21.31 locus indicate significant genetic differentiation driven by selection, while the extended ROH segments highlight the recent fixation of advantageous alleles. Reduced molecular diversity and low pairwise differences within the EAS further underscore the impact of selective sweeps on genetic variation. The significantly negative Tajima’s D value provides direct evidence of non-neutral evolution, consistent with recent positive selection. Linkage disequilibrium decay analysis revealed that East Asians have a more gradual decay compared to other populations, indicative of extended haplotype structures, suggesting that either reduced recombination rates or strong, recent positive selection has maintained these haplotypes. XP-EHH analysis identified SNPs with significantly high positive scores in EAS, particularly in the genomic region that overlaps with the LZTFL1 gene. This gene has previously been linked to COVID-19 severity, suggesting a potential role in pathogen-driven adaptation. Relate’s marginal-tree reconstruction pinpoints LZTFL1 variant that swept to a high frequency in EAS, most likely in response to ancient Corona or related virus ∼ 20-25 Kya. Pathway analyses implicate BBSome-mediated cargo targeting and cilium assembly, critical for mucociliary clearance. The GENOMIC COVID-19 GWAS implicates 3p21.31 in severe COVID-19 in Europeans and South Asians, with the LZTFL1 variant showing a strong protective effect; the non-significant but negative effect in EAS suggests population-specific selection at this locus, possibly driven by an ancient coronavirus-like epidemic. Together, these findings suggest that the 3p21.31 locus plays a critical role in population-specific adaptation in EAS, likely in response to environmental or pathogenic challenges.

## Materials and Methods

### Genotype data and quality control

All analyses used chromosome 3 genotype data from the 1000 Genomes Project Phase 3 (GRCh37). To ensure the integrity of the dataset, genotype data were subjected to rigorous quality control (QC) in PLINK v1.9 (Chang et al. 2015). Sample-level QC was performed to remove individuals with a missing genotype rate > 0.01 (--mind 0.01) and variant-level QC removed markers with a missing call rate > 0.01 (--geno 0.01). Rare variants with minor allele frequency (MAF) < 0.01 were excluded (--maf 0.01) for downstream analyses.

### PBS analysis

We employed the PBS to investigate the selection signals. PBS quantifies lineage-specific selection by comparing allele frequency differentiation between the three populations and leveraging pairwise F_ST_ values (Weir and Cockerham 1984) to estimate divergence. We used the EUR and SAS as the reference population and East Asians (EAS) as the target population. The initial Fst for each combination were computed using vcftools (Danecek et al. 2011), and later it was used for PBS estimations.

The PBS was calculated for each locus using the following formula:

1. Branch length (T) for each population pair:

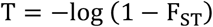
2. PBS for a specific branch:

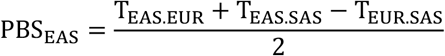

Where T_EAS.EUR_, T_EAS.SAS_, and T_EUR.SAS_ represent transformed pairwise F_ST_ values between population pairs based on the equation T = −log(1 − FST). A stringent threshold was set at the 99.99th percentile of the PBS distribution to identify loci potentially under selection. This approach focuses on loci with extreme values, which indicate strong selection in specific branches. All statistical analyses, including PBS calculations and outlier detection, were performed using custom R and Python scripts. The visualisations of the PBS distribution and significant outliers were generated using R (Ihaka and Gentleman 1996).

### ROH analysis

We utilise the 1000 Genomes database to estimate ROH (Durbin et al. 2010). PLINK 1.9 (Chang et al. 2015) was used for data management. The ROH for each population was calculated using PLINK 1.9 (Chang et al. 2015). We used the --homozyg function to perform the analysis. For the calculations, we used a 100 kb window size with a minimum of 50 single-nucleotide polymorphisms (SNPs) per window, allowing one heterozygous and five missing calls per window. The designated window sequentially scanned each individual and estimated the proportion in a homozygous window for every SNP.

### Molecular indices, average pairwise and Tajima D analysis

A Perl script was used to convert the 3p21.31 locus Plink file to fasta (ped to IUPAC). DNAsp (ver. 6) was used for phasing and generating the Arlequin input file (Rozas et al. 2017). Arlequin 3.5 (Excoffier and Lischer 2010) was used to calculate the Molecular indices, Tajima D, and average pairwise and Nei’s genetic distance, which were then plotted on a graph using R (Ihaka and Gentleman 1996).

### LD Decay analysis

To investigate the decay of LD, we utilised PopLDdecay v3.43 (Zhang et al. 2019). The analysis was performed using high-quality SNPs from our dataset, ensuring the exclusion of low-quality variants (e.g., missing call rate >1% and minor allele frequency <0.01). Pairwise LD (measured as r^2^) was calculated across genomic loci, and the decay of LD with increasing physical distance was analysed. The results were visualised using the Perl script to assess differences in LD decay patterns among populations to infer potential selection signatures at this critical genomic region associated with COVID-19 severity.

### XP-EHH analysis

The XP-EHH analysis was performed using Selscan (Szpiech 2024) to detect positive selection at the 3p21.31 locus across different populations. High-quality genotype data were used as input (missing call rate >1% and minor allele frequency <0.05), and XP-EHH scores were computed by comparing haplotype structures between populations (EUR Vs EAS). The analysis focused on identifying genomic regions with extended haplotype homozygosity in one population relative to another. The results were normalised and visualised to highlight candidate loci under selection.

### Marginal Genealogical Tree Reconstruction, Detection of LD and Pathway analysis

Relate was used to estimate genealogies that adapt to changes in local ancestry caused by recombination to plot marginal genealogical trees (Speidel et al. 2019) for the candidate SNP identified through XP-EHH. We further used LDlink (Machiela and Chanock 2015) to identify proxy variants for our target SNP within the populations identified by our XP-EHH analysis. We then performed pathway annotations on these proxy variants using the SNPnexus (Oscanoa et al. 2020).

### Ancestry stratified GWAS Investigation of 3p21.31

We analysed publicly available summary statistics from the GENOMIC COVID-19 GWAS meta-analysis (Pairo-Castineira et al. 2023) to investigate genetic variation at the 3p21.31 locus. The SNP rs9871972, located within the LZTFL1 gene and residing in a strong linkage disequilibrium block, was selected for population-specific analysis. Effect sizes (β coefficients) and p-values were extracted for EUR, SAS, and EAS populations. Statistical associations with severe COVID-19 were assessed using the reported GWAS summary statistics, and a Manhattan plot was generated to visualise genome-wide significance patterns across the locus. Analyses were conducted independently for each population to account for potential differences in allele frequency and haplotype structure.

## Acknowledgments

RKP is supported by the ICMR SRF fellowship (2021-15707). SD is supported by the CSIR-JRF fellowship. GC is supported by ICMR ad hoc grants (2021-6389), (2021-11289) and BHU IoE incentive grant BHU (6031).

## Author Contributions

RKP and GC conceived and designed this study. RKP and SD analysed the data. RKP, SD and GC wrote the manuscript. All authors contributed to the article and approved the submitted version.

## Data availability

1000 Genome phase 3 data was downloaded from: https://ftp.1000genomes.ebi.ac.uk/vol1/ftp/

Using the inferred genealogies, downloaded from: https://www.dropbox.com/scl/fo/7wl6ubqkp4u317u1wz4cc/h?dl=0&e=1, and the genealogy at rs1058961 was plotted across archaic and modern human populations using the TreeView module of Relate.

COVID-19 GWAS summary statistics for ancestry-specific meta-analysis from GenOMICC study Release 3.1 were downloaded from: https://genomicc.org/data/r3.1/

